# Apical caspase reporters uncover unknown stages of apoptosis and enable ready visualization of undead cells

**DOI:** 10.1101/419630

**Authors:** Luis Alberto Baena-Lopez, Lewis Arthurton, Marcus Bischoff, Jean-Paul Vincent, Cyrille Alexandre, Reuben McGregor

## Abstract

The caspase-mediated regulation of many cellular processes, including apoptosis, justifies the substantial interest in understanding all of the biological features of these enzymes. To complement functional assays, it is critical to identify caspase-activating cells in live tissues. Our work describes new caspase-reporters that, for the first time, provide direct information concerning the initial steps of the caspase activation cascade in *Drosophila* tissues. One of our caspase-sensors has capitalized on the rapid subcellular localization change of a fluorescent marker to uncover novel cellular apoptotic events. These refer to the actin-mediated positioning of the nucleus before cell delamination. The other construct has benefited from a caspase-induced nuclear translocation of a QF transcription factor. This feature enables the genetic manipulation of caspase-activating cells, whilst showing the spatio-temporal patterns of apical caspase activation. Collectively, our sensors offer new experimental opportunities that are already illuminating unknown aspects of caspase-dependent processes in apoptotic and non-apoptotic cellular scenarios.

**Summary statement:** We describe a novel set of caspase sensors that directly detect early caspase activation. The exclusive features of our reporters uncovered unknown stages of apoptosis and properties of caspase-activating cells.

## INTRODUCTION

The cysteine-dependent aspartate proteases, commonly known as caspases, are the major regulators of apoptosis, but also decisively modulate other essential biological functions independent of apoptosis (e.g. cell proliferation, cell differentiation and cell migration) (Aram et al., 2017; Baena-Lopez et al., 2017; Burgon and Megeney, 2017; Ellis and Horvitz, 1986; Fogarty and Bergmann, 2017; Hollville and Deshmukh, 2017; McIlwain et al., 2013; Miura, 2012; Mukherjee and Williams, 2017; Perez-Garijo, 2017; Songane et al., 2018). Accordingly, caspase malfunction in either apoptotic or non-apoptotic cellular scenarios often leads to disease initiation and progression (Aram et al., 2017; Baena-Lopez et al., 2017; Burgon and Megeney, 2017; McIlwain et al., 2013; Miura, 2012; Mukherjee and Williams, 2017; Perez-Garijo, 2017). As a first step towards characterising all of the caspase related biological functions within complex cellular assemblies, it is important to devise methods that efficiently identify caspase-activating cells in live tissues. However, the repertoire of caspase sensors with such properties is currently limited (Bardet et al., 2008; Ding et al., 2016; Florentin and Arama, 2012; Schott et al., 2017; Takemoto et al., 2003; Tang et al., 2015; To et al., 2015).

Apoptotic caspases have been grossly classified as either initiator/apical or executioner/effector depending on their early or late activation during the apoptosis (Baena-Lopez et al., 2017; Ramirez and Salvesen, 2018). The genetically encoded caspase-sensors described thus far, are based on short caspase-cleavage sites (DEVD or DQVD), specifically recognised by effector caspases (Bardet et al., 2008; Ding et al., 2016; Florentin and Arama, 2012; Schott et al., 2017; Takemoto et al., 2003; Tang et al., 2015; To et al., 2015). In order to make these reporters compatible with live imaging techniques, they have often incorporated different fluorophores at both ends of the caspase-recognition site (Bardet et al., 2008; Florentin and Arama, 2012; Schott et al., 2017; Takemoto et al., 2003; To et al., 2015). Some of these sensors have exploited changes in the subcellular localization of fluorescent proteins to visualize caspase activation (Bardet et al., 2008), whilst others have relied on split fluorophores that exclusively fluoresce upon caspase-mediated cleavage (Schott et al., 2017; To et al., 2015). Although these methods are undoubtedly powerful, even in non-lethal scenarios (Kanuka et al., 2005), they share some limitations. They cannot provide a temporal perspective of caspase activation over long periods of time; nor do they enable straightforward genetic manipulation of caspase-activating cells. Moreover, their activation requires the enzymatic activity of effector caspases, and therefore they are not functional in biological contexts without the participation of the entire caspase pathway; a situation frequently observed in non-apoptotic scenarios (Kondo et al., 2006; Napoletano et al., 2017; Ouyang et al., 2011; Wells et al., 2006). Some of these issues have been partially overcome by two recent constructs that have incorporated a CD8 membrane retention domain and a transcriptional activator (Gal4) flanking the caspase-cleavage motif (Ding et al., 2016; Tang et al., 2015). However, these reporters still rely on an effector caspase cleavage motif (DQVD), and the inclusion of a Gal4 fragment impedes their usage in combination with pre-existing Gal4 lines.

Here, we describe a new set of highly sensitive caspase-reporters that overcome all the aforementioned shortcomings, by incorporating an enzymatically dead but still cleavable template of the effector caspase Drice. This configuration ensures direct excision by apical caspases, while preventing their ability to trigger apoptosis. Our reporters also include additional features that have proven useful in unearthing new nuclear movements in pre-apoptotic cells as well as unknown biological properties of the caspase-activating cells in different *Drosophila* tissues.

## RESULTS

### Rational design of a novel Drice-based sensor (DBS)

Drice is fully activated by two sequential steps of enzymatic processing, with the first cleavage step being mediated by apical caspases (mainly by Dronc, Figs. S1A and 1A) (Lannan et al., 2007). Upon this first cleavage, Drice is split into two subunits (large and short), which remain strongly associated to form the active protease (Fig. S1A) (Lannan et al., 2007). We capitalised on this processing step to devise a reporter of apical caspase activation, which will be hereafter referred to as the Drice-based sensor (DBS). As a foundation for the construct, we used an enzymatically inactive but still cleavable version of Drice; Drice^C211A^ (Fig. 1A) (Lannan et al., 2007). This construct configuration does not compromise the apical caspase-mediated excision events but prevents undesirable activation of apoptosis (Lannan et al., 2007). We then appended the transmembrane domain of CD8 at the N-terminus and a Histone2Av-GFP moiety to the C-terminus (Figure 1A). Following this design, we created two versions of DBS. One version included a full-length form of Drice (CD8-Drice[C211A]-full length-Histone-GFP; DBS-FL), while the other contained a truncated template that only retained sixteen aminoacids downstream of the Dronc cleavage site (CD8-Drice[C211A]-short-Histone-GFP; DBS-S) that hypothetically prevents the interaction between the large and the small subunit of Drice (Figure 1A). Both constructs were subcloned downstream of the ubiquitous *tubulin* promoter to assess their activity in S2 *Drosophila* cells and transgenic flies. In the absence of caspase activation, as expected, both constructs are anchored to the cellular membranes outside the nucleus (notice the predominant membrane GFP signal in 91% of the transfected cells; n=63 cells; Figs. 1B, S1B and S1C). However, 90 minutes after inducing cell death with UV light, the excision of the small Drice subunit in the DBS-S construct allowed the translocation of the Histone-GFP moiety into the nucleus within most of the transfected cells (78% (n=77); Fig. 1B and S1B). As confirmation that DBS-S faithfully reports on caspase activity, we found that the nuclear localization of the Histone-GFP fragment was correlated with caspase-3 immunoreactivity (Fig. 1B). Additional evidence to discard unspecific cleavage of our sensor outside of the Drice template was obtained by observing that the fluorescent signal of DBS-FL remained attached to the membranes even in apoptotic conditions, highly likely due to the strong interaction between the large and small subunits of Drice (Fig. S1C). Equivalent observations were made expressing the sensor under the regulation of a different promoter (*actin* promoter; Fig. S1D). These results suggest that DBS-S is able to report on caspase activation in apoptotic cells, and there is no inadvertent or unspecific cleavage of DBS-S template without apoptotic stimuli.

**Figure 1.**
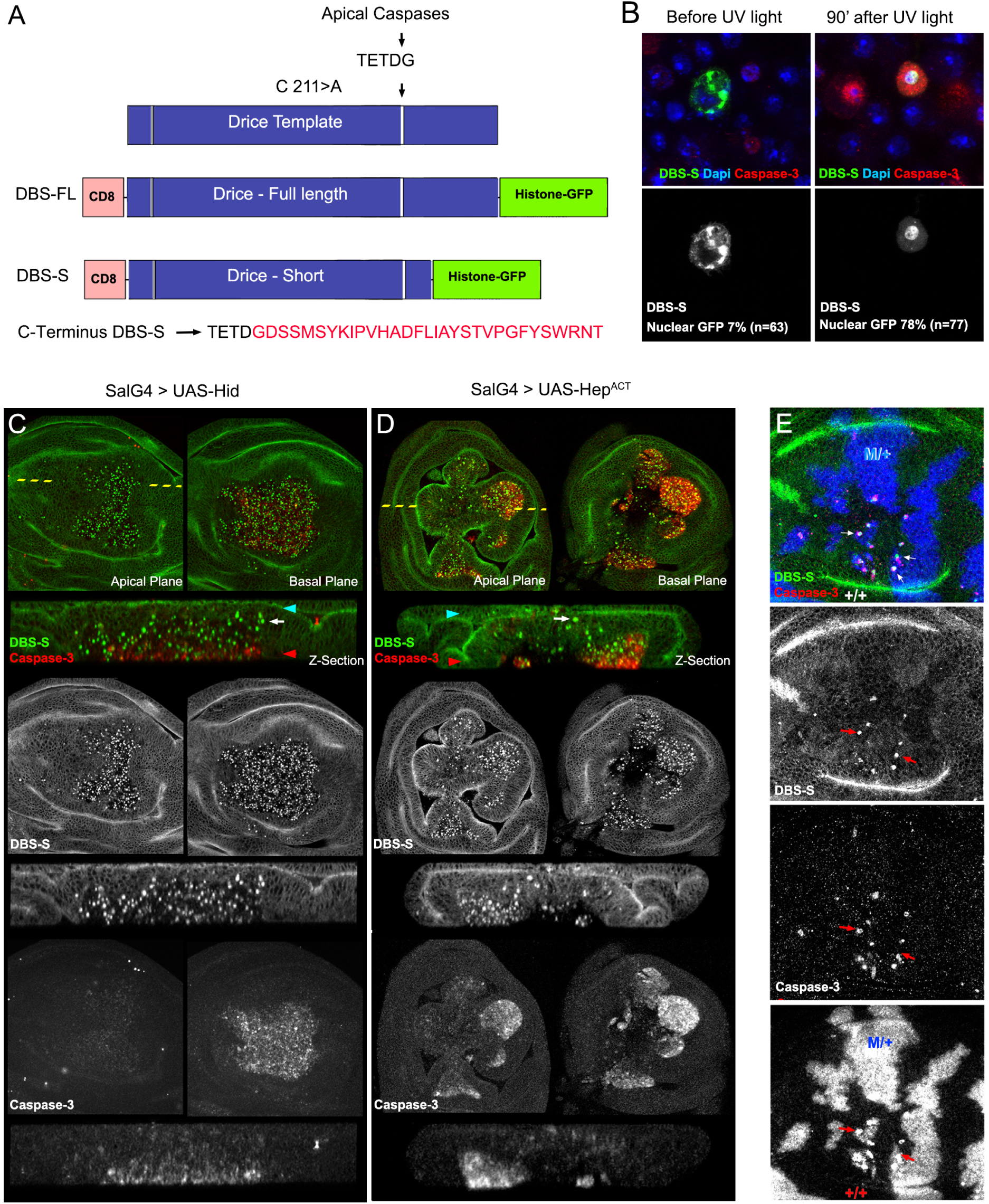
Rational design and validation of Drice-based sensor (DBS-S) **A)** Schematic diagram of Drice template indicating the mutation introduced in the catalytic residue (C 211> A; Cysteine to Alanine substitution) and the apical caspase cleavage site (TETD/G). Rational design of constructs generated from Drice template (Drice based sensor-full length-histone GFP, DBS-FL and Drice based sensor-short-histone GFP, DBS-S). Amino acid sequence kept in DBS-S after the apical cleavage site (shown in red). **B)** Functional characterization of DBS-S in S2 cells before and after UV light exposure DBS-S, caspase-3 and Dapi are shown in the green/grey, red and blue channels, respectively**. C and D)** Functional characterization of DBS-S in *Drosophila* wing discs, expressing the pro-apoptotic protein Hid (C) or Hemipterous activated (D) under the regulation of *spalt*-Gal4 (DBS-S > green and caspase-3 > red). *spalt*-Gal4 is expressed in the central region of the wing pouch where the nuclei are positively labelled with the sensor. Shown in the panels are XY images of apical and basal focal planes of the wing discs and cross-sections of the epithelia (Z-sections). The discontinuous yellow line indicates the location of the Z-sections in the XY images. Blue and red arrowheads indicate apical and basal regions of the epithelium in the Z-sections, respectively. White arrows point cells apically located activating DBS-S (i.e. nuclear GFP). **E)** Physiological cell death triggered by cell competition mechanisms. Genetic Minute heterozygous mosaics (M/+ cells > blue) showing the colocalization of nuclear DBS-S signal (green) and caspase-3 immunoreactivity (red) in presumptive apoptotic cells (red arrows). Please see full description of the experimental genotypes displayed in all the figures in MM.

### DBS-S is an early reporter of cell death in Drosophila cells

To investigate the activity of DBS-S in *Drosophila* tissues, we generated transgenic flies expressing the reporter under the regulation of ubiquitous promoters (*tubulin* or *actin*). The resulting transgenic flies were developmentally normal, fertile and morphologically indistinguishable from their wild-type siblings. These observations indicate that DBS-S expression did not compromise developmental processes that require either apoptotic cell death or non-apoptotic caspase activation (e.g. dorsal closure (Bischoff and Cseresnyes, 2009; Levayer et al., 2016; Marinari et al., 2012; Ninov and Martin-Blanco, 2007) or differentiation of sensory organs (Kanuka et al., 2005)). Since wild-type imaginal discs have a low rate of spontaneous apoptosis in standard laboratory conditions, the nuclear labelling with DBS-S was extremely rare in this tissue (Fig. S1E). However, many nuclei were positively labelled in response to genetically induced apoptosis (Figs. 1C, 1D and S2A), environmental apoptotic stimuli (Figs. 2, S2B and S2C), or physiologically triggered cell death (Figs. 1E and S2D) (Moreno et al., 2002). In pro-apoptotic conditions, cells positively marked with the sensor often had caspase-3 immunoreactivity (Figs. 1C-1E) and were located at the basal surface of the epithelium (Figs. 1C and 1D). Additionally, we observed GFP-positive nuclei apically located in cells that had either low or no caspase-3 immunoreactivity (Figs. 1C and 1D). Similar results were obtained utilising all of the different transgenic lines expressing the construct (Figs. 1 and S2). Western blot experiments also confirm the cleavage of the sensor following the expected pattern upon apoptosis induction (Fig. S1F). These observations suggest that DBS-S is able to track the early stages of apoptosis, before compromised cells initiate the delamination process (Gudipaty and Rosenblatt, 2017), while confirming the specific cleavage of the sensor in response to apoptotic stimuli in either induced or physiological situations. Next, we investigated the performance of DBS-S in non-apoptotic scenarios, analysing its activation within the proneural clusters of the wing discs (Kanuka et al., 2005). However, no nuclear fluorescent signal was observed during the course of these experiments (not shown). These results suggest that DBS-S mainly detects the high levels of caspase activation that accompanies apoptotic cells.

**Figure 2.**
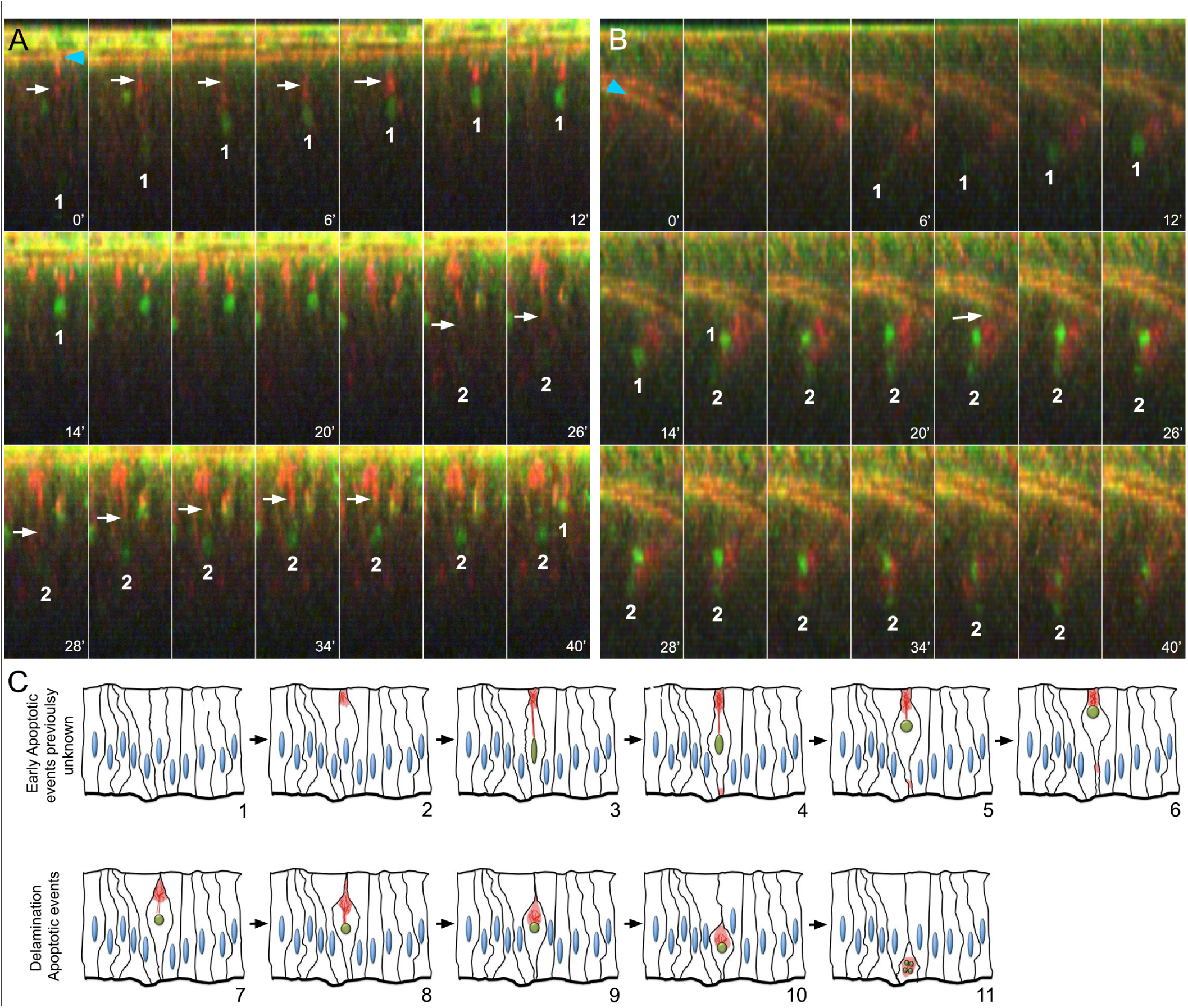
Live imaging of imaginal discs upon irradiation. **A)** Time-lapse of wing imaginal discs *ex vivo* (cross-sections) after irradiation (time is indicated in minutes; DBS-S is show in green; LifeAct-Ruby shows actin in red). Notice the progressive accumulation of GFP signal in the nuclei of cell 1 and 2, the changes in their shape (roundness increase) and the movement towards the apical cell cortex in A. White arrows indicate the acting bundles connecting the apical cell cortex with the nuclei; notice the contraction of the actin bundle over time. The recording started 90 minutes after irradiation and lasted for 40 minutes. Snapshots were acquired every 2 minutes. **B)** Delamination process of apoptotic cells; white arrow indicates apical cell detachment before delamination. Blue arrowheads indicate the normal accumulation of actin in the apical adherent cell junctions. Notice that actin remains accumulated around the nuclei during the movement towards the basal surface of the epithelium. Image acquisition conditions are described in A. **C)** Schematic representation of early apoptotic events newly identified by DBS-S and LifeAct-Ruby in irradiated wing discs (1-6), as well as delamination events previously described during apoptosis (7-11). The activation of the reporter is indicated by the green colour of the nuclei, while actin network is represented in red.

### Apical nuclear migration in pre-apoptotic cells uncovered by DBS-S

After completing the basic characterisation of DBS-S in fixed samples, we analysed the performance of the sensor in live tissues. DBS-S readily identified apoptotic cells in imaginal discs filmed *ex-vivo* upon irradiation (Movie S1). Strikingly, we observed that most of the GFP positive nuclei aligned to the apical surface soon after irradiation (Figs. 2A, Figure S2C, Movie S1, S2 and S3). This movement was tightly correlated with the apical accumulation of actin (Fig. 2A, Movie S2 and S3). Indeed, the contraction of actin bundles appeared to pull the nuclei up towards the apical cell cortex, whilst the nuclei kept accumulating the GFP signal (Figs. 2A and S2C, Movie S2, Movie S3). GFP-positive nuclei also change their shape during the apical migration, progressively becoming more rounded (Figs. 2A and S2C, Movie S2, Movie S3). Subsequently, during the delamination process, the nuclei were pushed towards the basal side of the epithelium (Fig. 2B, Movie S4) along with the actin-enriched structure (Figs. 2B and 2C, Movie S4). During most of the delamination process, the nuclei remained intact (Figure S1E), therefore nuclear fragmentation appears to be a characteristic event of apoptotic cells in the final stages of delamination. A similar association between nuclei and polymerized actin was also noticed in apoptotic S2 cells (Fig. S1D). These results confirmed that DBS-S is an early sensor of apoptosis, which in turn has allowed us to uncover unknown behaviours of pre-apoptotic nuclei (Fig. 2C and Movie S5). DBS-S was also an effective marker of apoptotic cells in other tissues beyond the wing discs. Thus, DBS-S marked characteristic apoptotic larval epidermal cells during pupal stages (Fig. S2D, Movie S6) (Ninov et al., 2007) and apoptotic histoblasts (Movie S7) of which 3% die during metamorphosis, (Bischoff and Cseresnyes, 2009)). Exceptionally in embryos, either physiological or experimental induction of apoptosis (with Ci-gal4 and UAS-reaper) did not lead to the activation of the DBS-S sensor (not shown). However, this uncertain disparity does not undermine the power of the DBS-S to identify apoptotic cells in most situations.

### Sensitivity of DBS-S to apical caspases and P35

Although *dronc* is considered the main apical caspase in *Drosophila*, other initiator caspases (*dredd* and *strica*) could also contribute to Drice activation. To address whether Dronc is the sole activator of DBS-S, we genetically induced apoptosis in a *dronc* null mutant background (*dronc*^I29^) by ectopically activating JNK-signalling. Imaginal discs of such genetic condition are phenotypically hyperplastic and still show a limited number of caspase-3 positive cells (Perez-Garijo et al., 2009). Confirming previous results, caspase-3 immunoreactive cells were also marked with DBS-S (Figs. 3A and 3B). These findings suggest that other apical caspases could residually mediate the cleavage of Drice and therefore DBS-S template without Dronc in pro-apoptotic conditions (see discussion).

**Figure 3.**
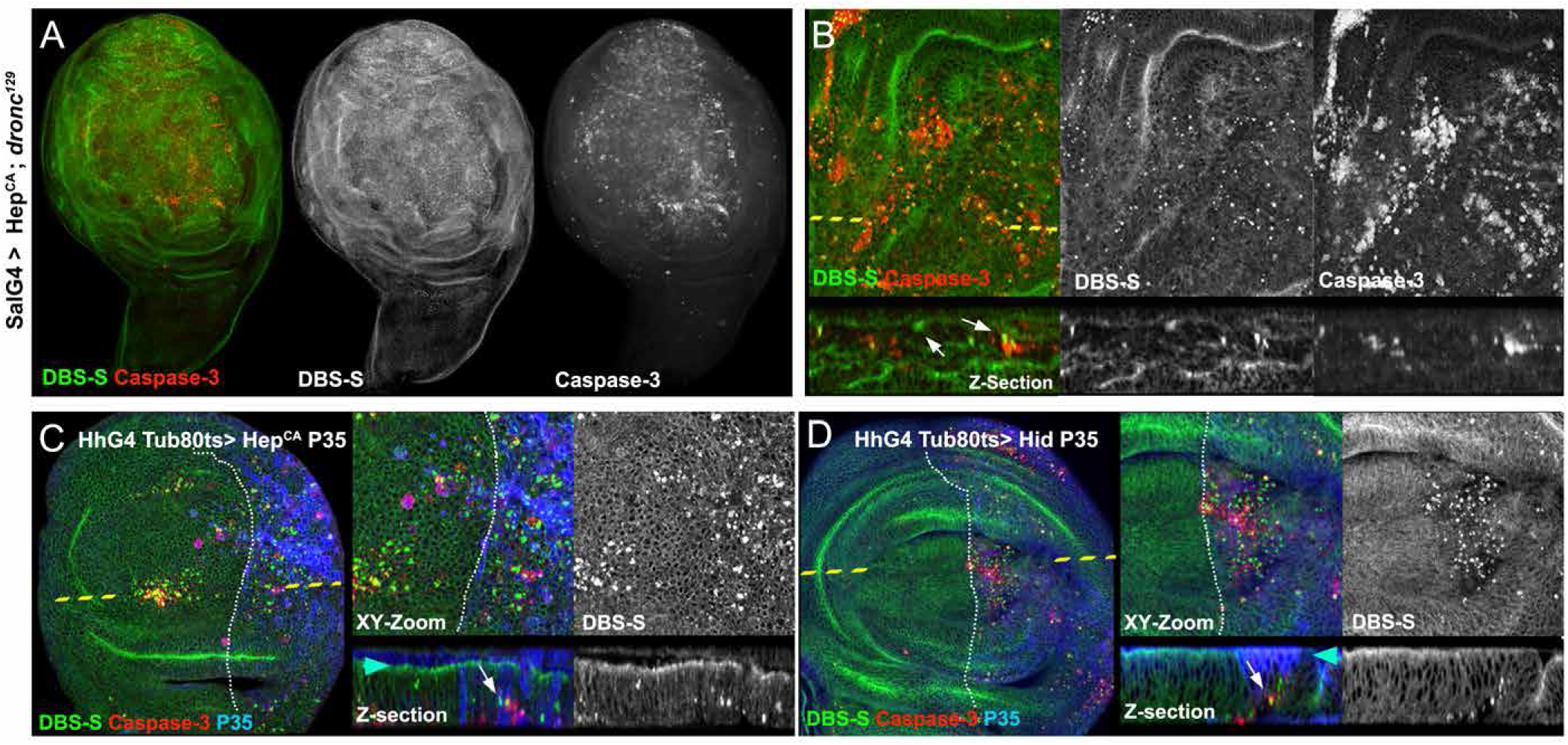
Behaviour of the DBS-S in compromised cells that cannot complete the apoptosis programme. **A)** Wing cells mutant for *dronc* that ectopically express an activated form of Hemipterous under the control of *spalt*-Gal4 (DBS-S > green, caspase-3 > red). **B)** Detail from A. Notice the presence of cells positively labelled with DBS-S (nuclear signal of the reporter > green, white arrows) and caspase-3 immunoreactivity (red). **C)** Identification and labelling of undead cells in wing imaginal discs expressing simultaneously Hemipterous activated and P35 under the control of Hedgehog-Gal4. XY images, high magnifications and cross-sections of wing epithelium (Z-sections) are shown in C. DBS-S is shown in green, caspase-3 staining in red and P35 immunostaining in blue. The anterior (left side) and posterior (right side) compartments are separated by the discontinuous white line. The discontinuous yellow line indicates the location of the Z-section in the XY image. White arrows indicate delaminated cells positively labelled with DBS-S (green). Blue and red arrowheads indicate the apical and basal parts of the wing epithelium, respectively. **D)** Identification and labelling of undead cells in wing imaginal discs simultaneously expressing Hid and P35 under the control of Hedgehog-Gal4. Figure labelling is equivalent to the description of panel (C).

Next, we investigated the impact of the effector caspase inhibitor P35 on the DBS-S (Hay et al., 1994). P35 covalently binds to the catalytic residue of effector caspases compromising their activity to cleave substrates (Hay et al., 1994; LaCount et al., 2000; Lannan et al., 2007). Accordingly, it also blocks the activation of effector caspase reporters (Bardet et al., 2008; Ding et al., 2016; Florentin and Arama, 2012; Schott et al., 2017; Takemoto et al., 2003; Tang et al., 2015; To et al., 2015). In these experiments, we used both Hemipterous (an activator of JNK-signalling) and Hid, as inducers of the apoptotic program. Since P35 acts downstream of *dronc*, it did not prevent the nuclear import of GFP (Figs. 3C and 3D). As expected, this indicates that DBS-S is insensitive to inhibitors of the effector caspases. Additionally, it supports the notion that DBS-S cleavage is specifically mediated by apical caspases (see next section). Importantly, this feature brings up the possibility of identifying caspase-dependent processes with our sensor that do not require the entire caspase cascade (e.g. tracking of the so-called “undead cells” (Perez-Garijo et al., 2005); Figs. 3C and 3D).

### DBS-S-QF: a temporal caspase reporter that allows the genetic manipulation of caspase-activating cells

To increase the sensitivity and broaden the usefulness of our sensor, we decided to replace the Histone-GFP-encoding fragment in DBS-S by the transcriptional activator QF (Potter et al., 2010) (DBS-S-QF; Fig. S3A). As previously shown by DBS-S transgenic flies, the constitutive expression of DBS-S-QF did not generate morphological or developmental defects, thus discarding either toxic or inadvertent side effects of the construct. Likewise the DBS-S-QF reporter retained most of the features of DBS-S, such as the labelling of apoptotic cells (Figs. 4A, 4B, S3B and S3C). Although this feature is strongly linked to the cellular marker activated by DBS-S-QF (whereas P53-expressing cells of the wing can activate the QUAS-lacZ transgene, failed to show activation of QUAS-Tomato-HA; compare Figs. S3B with S3D), as well as the tissue analysed (whereas P53-expressing cells activated the QUAS-Tomato-HA transgene in the eye imaginal discs, failed to show expression of the same transgene in the wing discs; compare Figs. S3C with S3D; see discussion). Importantly, the basal activation of DBS-S-QF also disappears by eliminating the expression of *dronc* (Figure 4C and 4D). Conversely, P35 expression do not prevent the cleavage of DBS-S-QF, allowing “undead” cells to be tracked (Fig. 4F, notice in this instance the activation of QUAS-tomato in comparison to controls 4E, see discussion). By virtue of releasing the transcription activator QF, DBS-S-QF is also sensitive to detect caspase activation in non-apoptotic contexts (SOPs; Fig. S3E). These results collectively confirm the apical caspase-dependent cleavage of the DBS template in apoptotic and physiological conditions. Additionally, the QF release upon caspase cleavage from DBS-S-QF enables the genetic manipulation of caspase-activating cells in combination with the Gal4/UAS system (Figs. 4C-F, S3E-3G). Taking advantage of this feature, we replicated previously described patterns of caspase activation in wing imaginal discs (Ding et al., 2016; Tang et al., 2015) (Figs. 4G and 4H); but we also observed striking differences in other tissues such as the posterior midgut of adult flies. Whereas effector caspase-based sensors were exclusively activated in enterocytes throughout the posterior midgut (Ding et al., 2016; Tang et al., 2015), DBS-S-QF was also activated within the intestinal progenitor cells (intestinal stem cells and enteroblasts; Figs. 5A, 5B and Figure S4A). Additionally, although this activation was initially localised in the posterior region of the midgut (R4c region (Buchon et al., 2013)), the activation spread anteriorly over time (R4a and R4b; Figs. 5C-E and S4A). Importantly, sensor-positive cells did not show signs of apoptosis and remained in the epithelium (Fig. S4B). Beyond confirming the widespread non-lethal caspase activation (Ding et al., 2016; Tang et al., 2015), our results suggest the existence of stereotyped patterns of apical caspase activation, likely to be regulated by either developmental or environmental cues in the *Drosophila* gut.

**Figure 4.**
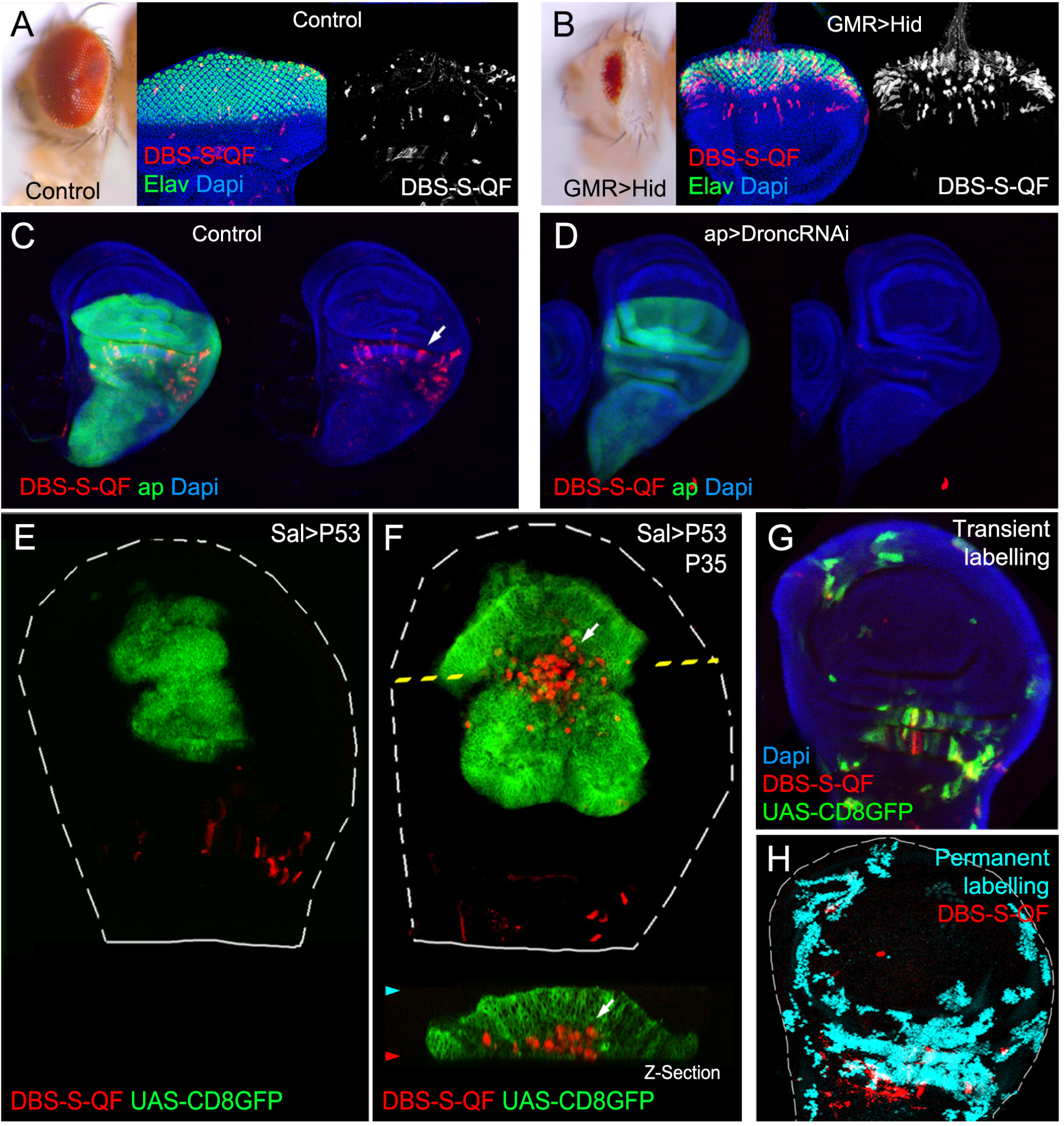
Functional characterization of DBS-S-QF sensor. **A and B)** Activation of DBS-S-QF reporter in either wild-type (A) or apoptotic eyes ectopically expressing Hid under the regulation of GMR promoter (B). Adult eyes and imaginal discs are shown on the left and right panels, respectively. Activation of DBS-S-QF is shown in red upon induction of a QUAS-tomato-HA transgene (anti-HA in red). Differentiated regions of the eye express the neuronal identity marker *elav* (green); nuclei are labelled with Dapi. **C)** Activation of DBS-S-QF (anti-HA in red, white arrow) in a wild type wing disc expressing CD8-GFP (UAS-CD8-GFP) under the regulation of *apterous*-Gal4 (green); the experiment was made at 29 °C. **D)** Activation of DBS-S-QF (anti-HA in red) in a wing disc expressing a RNA interference against *dronc* (UAS-Dronc-RNAi) and CD8-GFP under the regulation of *apterous*-Gal4 (green); the experiment was made at 29 °C **E)** Overexpression of P53 and CD8-GFP under the regulation of *spalt*-Gal4 (green); red channel shows the DBS-S-QF activation (QUAS-tomato-HA, anti-HA in red). Notice the absence of Tomato-expressing cells in the GFP expressing cells. The discontinuous white line outlines the wing disc. **F)** Co-expression of P53, P35 and CD8-GFP in the *spalt*-Gal4 expression domain (green); red channel shows the DBS-S-QF activation (QUAS-tomato-HA, anti-HA in red); white arrow in XY-image or Z-section indicates the appearance of “undead cells” basally delaminated (undead cells refers to caspase-activating cells that cannot finalise the apoptosis programme due to the presence of P35 blocking the activation of the effector caspases). **G**) Temporal perspective of caspase-activating cells using transient fluorescent markers expressed under the control of DBS-S-QF; QUAS-tomato-HA (anti-HA in red) labels ongoing caspase activation; old caspase activation is shown in green (QUAS-Gal4 UAS-mCD8-GFP); recent-past caspase activation is indicated by the colocalization of both markers (yellow); Dapi labels nuclei. **H**) Lineage-tracing of caspase-activating cells in the wing discs obtained with DBS-S-QF (ongoing activation of DBS-S-QF is shown in red, while nuclear beta-gal staining shows in cyan the permanent labelling of caspase-activating cells).

**Figure 5.**
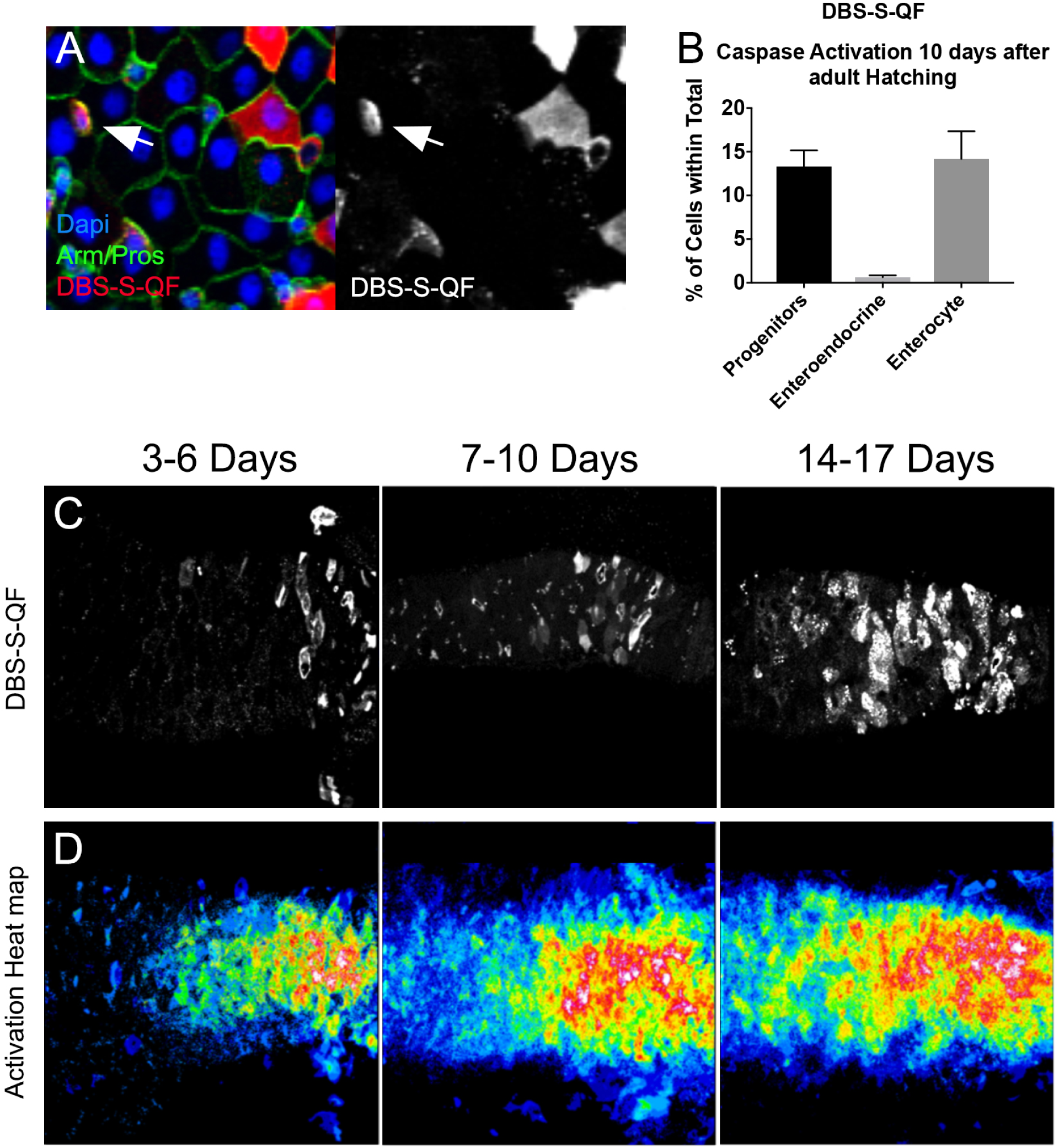
Functional characterization of DBS-S-QF in the posterior midgut of adult flies. **A**) Transient expression of DBS-S-QF in the midgut (QUAS-Tomato-HA, anti-HA in red); anti-Armadillo staining in green labels the cell membranes; nuclear anti-prospero staining in green labels enteroendocrine cells; Dapi staining marks nuclei). White arrows point intestinal progenitor cells (intestinal stem cells or enteroblasts) positively labelled by DBS-QF. **B**) Quantification of cell types transiently activating DBS-S-QF in the posterior midgut. Large nuclear cell size was the criteria to assign enterocyte cell identity. The criteria applied to identify gut progenitor cells (either intestinal stem cells or enteroblasts) was the small cell size and negative immunolabelling against Prospero. n=10. Error bars indicate standard error of the mean. **C and D**) DBS-S-QF activation in the gut at different time points during adulthood. Left panels show DBS-S-QF activation (QUAS-tomato-HA anti-HA in white), while right panels show the overlay of different guts at the same time point. Warmer areas (red) in the heat map indicate higher concentration of cells activating the sensor, while colder areas (blue) areas reflect less cells activating the reporter. DBS-S-QF: 3-6d n=13, 7-10d n = 25, 14-17d n = 24.

## DISCUSSION

The labelling of caspase-activating cells in live tissues is critical to identify caspase-dependent biological processes in apoptotic and non-apoptotic cellular scenarios. We have developed a new set of *Drosophila* caspase-sensors that provide a complementary view of caspase activation in comparison to currently available reporters, through directly monitoring the activation of apical caspases. Our sensors are initially retained at the cellular membrane through a CD8 transmembrane domain and exclusively, upon apical caspase activation (mainly Dronc), allow the release of either a Histone-GFP fragment or the transcriptional activator QF. These features have provided novel insights into caspase-activating cells in cellular scenarios that do not demand the activation of the entire caspase cascade.

### Novel apoptotic steps uncovered by DBS-S; apical nuclear migration of pre-apoptotic cells

Since the early 1970s, nuclear shape changes have been considered hallmarks of apoptosis, distinguishing this process from other programmes of cell death (such as necrosis or autophagy) (Kerr et al., 1972; Prokhorova et al., 2015). Indeed, nuclear shrinking and fragmentation are indicative of chromatin condensation and DNA degradation during apoptosis (Prokhorova et al., 2015). In *Drosophila*, these events occur relatively late and are tightly coordinated with the delamination process (Gudipaty and Rosenblatt, 2017; Kawamoto et al., 2016; Ohsawa et al., 2018) (Fig. 2C, delamination phases from step 7 onwards). Unsurprisingly, cell delamination and nuclear fragmentation are decisively influenced by caspase-mediated actin cytoskeleton rearrangements (Gudipaty and Rosenblatt, 2017; Kawamoto et al., 2016; Prokhorova et al., 2015). Our work now uncovers another actin-dependent event that occurs earlier during apoptosis. As the nuclear GFP signal from DBS-S accumulates; the nuclei are seen to undergo an apical migration and shape changes (an increase in roundness) (Figs. 2A, 2C and Movie S5). It is also tightly linked to the apical accumulation of actin and the subsequent contraction of an actin bundle connecting the apical cell cortex with the nucleus (Figs. 2A, 2C and Movie S5). These new apoptotic events precede the distinctive basal delamination movement of dying cells and, therefore, can be considered the earliest subcellular manifestation of apoptosis described to date in *Drosophila* cells. Interestingly, the uncovered apical migration of pre-apoptotic nuclei bears remarkable similarities with the interkinetic nuclear movement that normally occurs before mitosis (Meyer et al., 2011). However, the actin enrichment as well as the actin-mediated events appears to be localized in opposing subcellular domains in the two processes. Whereas the actin-dependent basal cellular contraction pushes nuclei towards the apical cortex during mitosis (Meyer et al., 2011), the retraction of apical actin bundles appears to pull nuclei upwards during apoptosis. Although it is still premature to establish the biological significance of this pre-apoptotic apical migration, we hypothesise that it could facilitate the rapid chromatin condensation and shutdown of gene expression, similar to the process occurring during mitosis (Zaret, 2014). Collectively, these events could inhibit essential cellular functions earlier than previously thought and before the DNA degradation is completed, thus ensuring the efficient elimination of apoptotic cells.

### Suitability of our sensors to detect caspase-related processes independent of effector caspases. Implication of caspases during cell delamination

In contrast to published sensors (Bardet et al., 2008; Ding et al., 2016; Florentin and Arama, 2012; Schott et al., 2017; Takemoto et al., 2003; Tang et al., 2015; To et al., 2015), our reporters are insensitive to inhibitors of the effector caspases. This feature has enabled for the first time the specific detection of apical caspase activation in cells in which the completion of the apoptotic program was experimentally prevented (e.g. “undead cells”). Indeed, visualization of “undead cells” with DBS-S-QF has allowed us to demonstrate that even though these cells do not undergo apoptosis, they can still delaminate (Figs. 3C, 3D and 4F). This suggests that the threshold of caspase activation required for cell delamination is lower than that for organelle destruction during apoptosis. It also suggests that cell delamination could be exclusively driven by apical caspases. This observation is however very puzzling since it is known that effector caspases are essential for the physiological cell delamination in the thorax (Levayer et al., 2016; Marinari et al., 2012). One possibility is that apical caspase activation is much stronger in our experimental conditions due to the overexpression of pro-apoptotic factors and consequently, sufficient for cell delamination. Alternatively, the process of caspase-mediated cellular delamination could be different depending on the tissue architecture.

### The faster you proliferate, the quicker you disappear

DBS-S takes advantage of rapid changes in the subcellular localization of the GFP signal to label caspase-activating cells, whereas its equivalent QF version demands a more complex sequence of events (e.g. QF translocation to the nucleus, transcriptional activation of the reporter gene and protein maturation). These additional steps confer higher sensitivity to DBS-S-QF, but inherently delay the labelling of apoptotic cells. Indeed, P53-apoptotic cells in the wing can be always detected with DBS-S but only with specific cellular markers activated by DBS-S-QF, presumably because dying cells are eliminated before they have opportunity to express the readout gene of choice (whereas P53-expressing cells of the wing can activate QUAS-lacZ, failed to show activation of QUAS-tomato-HA; compare Figs. S3B with S3D). Supporting this hypothesis, the co-expression of P35 and P53 prevented or slowed down the elimination of caspase-activating cells in the wing and consequently allowed the induction of the QUAS-tomato-HA signal under the regulation of DBS-S-QF (compare Figs. 4E and 4F). Interestingly, similar differences were observed between P53 fast-proliferating cells of the wing and non-proliferative cells behind the morphogenetic furrow of the eye (compare Figs. S3C and S3D). Therefore, we conclude that the ability of DBS-S-QF to label apoptotic cells largely depends on the disposal rate of apoptotic cells and the maker of choice induced by DBS-S-QF activation. Furthermore they illustrate the coordination between cell proliferation and cell destruction events previously hypothesised (Abrams and White, 2004).

### Apical vs effector caspase activation patterns

The spatio-temporal pattern of caspase activation revealed by our reporters and other “historical sensors” (Ding et al., 2016; Tang et al., 2015) have shown remarkable similarities and differences (Figure 5). The similarities are not surprising since the activation of previously described sensors demand the enzymatic activity of apical caspases. In this sense, our sensors would incorporate the activation patterns observed with effector caspase reporters. The discrepancies in the gut and potentially other tissues could be a consequence of apical caspase-mediated cleavage of our Drice-template. Importantly these disparities could be exploited to obtain specific information regarding biological processes that do not require the activation of effector caspases.

### Anastasis and non-apoptotic caspase functions

Our results confirm the widespread and non-apoptotic caspase activation previously noticed in many *Drosophila* organs (Ding et al., 2016; Tang et al., 2015). However, several factors suggest its likely relation with the regulation of specific cellular functions, instead of the anastasis phenomenon (cells activate caspases to trigger the apoptosis programme but eventually they are able to revert the course of events without completing the cell death (Ding et al., 2016; Tang et al., 2015)). Supporting this hypothesis, we find non-lethal caspase activation in a large number of cells in different tissues without signs of apoptosis (Figs. 4F and 5). Furthermore, the patterns of non-apoptotic caspase activation can be highly stereotyped, thus suggesting developmental control, as opposed to a response to random insults (Figs. 4H, 5, S4). Nevertheless, further functional studies are needed in years to come in order to establish the biological functions coupled to the non-apoptotic patterns of caspase activation in different organs.

In summary, we have described the functional characterization of novel apical caspase sensors that bring new experimental opportunities to uncover new biological features of caspase-activating cells in either apoptotic or non-apoptotic scenarios.

## MATHERIAL AND METHODS

### Fly Strains and experimental genotypes

All fly strains used are described at www.flybase.bio.indiana.edu unless otherwise indicated. All of our experiments were carried out at 25°C unless otherwise indicated. Full description of experimental genotypes:

*[w, Tubulin-DBS-S; UAS-hid (BL65403) /spaltEPV-Gal4 (Figure 1C)][w; UAS-Hep.Act (9306), Tubulin-DBS-S /spaltEPV-Gal4 (Figure 1C)] [yw heat shock-flipase/Tubulin-DBS-S; FRT40 w+/FRT40 Minute arm-LacZ (Figure 1E)] [w;;Actin mitoPlum-2A-LifeAct-Ruby-2A-DBS-S (Figure 2, S2B Movies S1-4)] [w;Tubulin-DBS-S (Figure S2D)] [w; UAS-Hep.Act (9306), Tubulin-DBS-S /spaltEPV-Gal4; dronc*^*129*^ *(Figure 3A and 3B)] [*w; *UAS-Hep.Act, Tubulin-DBS-S /TubG80*^*ts*^ *(from BL7019); Hedgehog-Gal4/UAS-P35 (BL5073) (Figure 3C)] [*w; *UAS-Hid, Tubulin-DBS-S /TubG80*^*ts*^ *(BL7019); Hedgehog-Gal4/UAS-P35 (Figure 3D)] [Actin DBS-S-QF, UAS-mCD8-GFP (from BL30118), QUAS-tomato-HA (from BL30118)/+ (Figure 4A)] [Actin DBS-S-QF, UAS-mCD8-GFP (from BL30118), QUAS-tomato-HA/+; GMR-Hid (from BL BL5251) (Figure 4B)] [Actin DBS-S-QF, UAS-mCD8-GFP, QUAS-tomato-HA; ap*^*md*^*544*^^- *Gal4 /+; (Figure 4C)] [Actin DBS-S-QF, UAS-mCD8-GFP, QUAS-tomato-HA; ap*^*md*^*544*^^*-Gal4 /UAS-Dronc-RNAi (gift from Pascal Meier); (Figure 4D)] [Actin DBS-S-QF, UAS-mCD8-GFP, QUAS-tomato-HA/+; spaltEPV-Gal4/GUS-Dp53 (BL6584) (Figure 4E)] [Actin DBS-S-QF, UAS-mCD8-GFP, QUAS-tomato-HA/+; spaltEPV-Gal4/GUS-Dp53; UAS-P35/+ (Figure 4F)] [Actin DBS-S-QF, UAS-mCD8-GFP, QUAS-tomato-HA/+;; QUAS-Gal4 (Figure 4G and S3I)] [Actin DBS-S-QF, UAS-mCD8-GFP, QUAS-tomato-HA/+; QUAS-FLPo (BL30126) /+; Actin5C FRT-stop-FRT lacZ-nls/+ (BL6355) (Figure 4H and S4B)][Actin DBS-S-QF, UAS-mCD8-GFP, QUAS-tomato-HA/+;; (Figure 5 and S4A)] [Tubulin-DBS-S/Cyo (Figure S1D)] [w; GUS-P53/spaltEPV-Gal4; Actin mitoPlum-2A-LifeAct-Ruby-2A-DBS-S/+ (Figure S2A)] [Actin DBS-S-QF, UAS-mCD8-GFP, QUAS-nucLacZ.7 (BL30006)/+; spaltEPV-Gal4/GUS-Dp53 (BL6584) (Figure S3B)] [Actin DBS-S-QF, UAS-mCD8-GFP, QUAS-tomato-HA/+;;GMR-Dp53 (BL8417) (Figure S3C)] [w, Actin DBS-S-QF, UAS-mCD8-GFP, QUAS-tomato-HA/+; GUS-P53/spaltEPV-Gal4 (Figure S3D)] [Actin DBS-S-QF, UAS-mCD8-GFP, QUAS-tomato-HA/+; neur*^*A*^*101*^^ *(BL4369)/+ (Figure S3E)] [Actin DBS-S-QF, UAS-mCD8-GFP, QUAS-tomato-HA/+;; QUAS-Gal4/UAS-Diap1 (Figure S3F and S3G)]*

### Molecular cloning and generation of new fly strains

All PCRs were performed with Q5 High-Fidelity polymerase from New England Biolabs (NEB, M0492L). Standard subcloning protocols and gene synthesis (Genewiz) were used to generate all the DNA constructs. New transgenic lines expressing our constructs were obtained by random P-element transgenesis and attP/attB PhiC31-mediated integration. Bestgene Inc. injected the DNA plasmids in *Drosophila* embryos in order to generate the new transgenic strains. Most of the plasmids designed for this study will be deposited at the DGRC, Indiana. Similarly, new fly strains generated will be deposited at the Bloomington Stock Centre. Whilst resources transfer is completed, sequences and reagents will be provided upon request.

#### Drice-Based Sensor-Histone-GFP constructs

A Bluescript vector containing DNA encoding for the N-terminal region (transmembrane domains) of mouse CD8 protein was used as a starting point. DNA encoding both the Drice mutant template (Drice C211A cDNA was a gift from Dr. P.D. Friesen) and the Histone-2AV GFP were amplified by PCR and cloned into ToPo-TA vectors. The Histone-2AV-GFP was then inserted at the unique BglII-Xba sites of the Bluescript vector lying after the CD8. The full-length and short Drice mutant templates where then inserted into the unique AgeI-BglII sites present between the CD8 and the Histone-2AV-GFP. Each construct was finally cloned as a NotI-XbaI fragment downstream of either *tubulin* or *actin* promoters included in a customized pCasper vector (this vector contains an attB-recombination site, a multicloning site and the fly selection marker white +). Random P-element insertions generated the DBS-S strains located in the II and III chromosomes, respectively. The integration of the construct on the X chromosome was obtained by using PhiC31-mediated recombination. The selected attP-landing site for generating the transgenic line was 9753.

#### DBS-S-QF reporter construct

To generate the QF version of DBS-S, the histone-2AV-GFP fragment was replaced by the open reading frame (ORF) of the transcriptional activator QF. We introduced a HA-motif at the N-terminus of the QF ORF and a BamHI restriction site via PCR. An extra SpeI site was also appended to the C-terminus of the PCR product. The Addgene plasmid (#46134) was used as template for obtaining the PCR fragment. The PCR product was then subcloned in the unique BglII-XbaI sites present in DBS-S-QF. The whole construct DBS-S-QF was finally transferred into an *actin* attb vector as a NotI-SalI fragment. The construct was integrated on the X chromosome using PhiC31-mediated recombination. The selected attP-landing site for generating the transgenic line was 9753.

#### Actin mitoPlum-2A-LifeactRuby-2A-DBS-S construct

We first generated a pMTV5 vector containing two repeated copies of the 2A peptide spaced conveniently with restriction sites (pMTV5 configuration NotI-BglII-2A-AvrII-ClaI-2A-XmaI-XbaI). Before the first 2A peptide we subcloned a plum fluorescent protein tagged with a mitochondrial localization signal. Gene synthesis and subsequent cloning as a NotI-BglII fragment was the strategy followed for the cloning of the Plum product. In between the two 2A peptides we cloned the PCR fragment encoding the lifeAct-Ruby protein (the template was a gift from Dr. E. Ober) using the unique sites AvrII-ClaI sites. After the second 2A peptide we cloned DBS-S as a fragment XmaI-XbaI. The whole construct Actin mitoPlum-2A-lifeActRuby-2A-DBS-S was finally subcloned into an *actin* attb vector as a NotI-XbaI fragment. The construct was integrated on the III chromosome using PhiC31-mediated recombination. The selected attP-landing site for generating the transgenic line was 9732.

#### QUAS-Gal4 construct

First we extracted the Gal4 ORF from pMTV5 Gal4 as a Not-AvrII fragment. Subsequently, the fragment was subcloned into a QUAS-attB vector containing a customized multicloning site. The construct was integrated on the III chromosome using PhiC31-mediated recombination. The selected attP-landing site for generating the transgenic line was 9748.

#### Primers description and sequences

Forward Histone2AV-GFP with BglII site: gatc**agatct**gctggcggtaaagcaggcaa

Reverse Histone2AV-GFP with XbaI site: gatc**tctaga**ttatttgtatagttcatccat

Forward Drice with AgeI site: gatc**accggt**gacgccactaacaatggagaatccgccg

Reverse Drice full length with BglII site: gatc**agatct**aacccgtccggctggtgccaactgcttgtcgc

Reverse Drice short with BglII site: gatc**agatct**gtgcactggaatcttgtagctcatcg

Forward HA-QF with BamHI site:

ttacac**ggatcc**aagctttacccatacgacgtccctgactatgcgcctccgggaattgggaattccaacatgccgcctaaacgca agacactcaatgc

Reverse QF polyA with SpeI site:

tttatata**actagt**ggatctctagaggtaccctcgagccgcggccgcggatctaaacgagtttttaagcaaactcactccctgaca ataaaaacgc

Forward LifeActRuby with AvrII site:

Tccggc**cctagg**aatgggtgtcgcagatttgatcaagaaattcgaaagcatctcaaaggaagaa

Reverse LifeActRuby with ClaI site:

tgccca**atcgat**agagcgcctgtgctatgtctgccctcagc

Forward CD8 with XmaI site

atgcgt**cccggg**atggcctcaccgttgacccgctttctgtcgc

Reverse Histone-2Av-GFP with XbaI site:

taaggg**tctaga**ttatttgtatagttcatccatgcc

### Mosaic Analysis

Females y w hsp70-flp; FRT40A M arm-lacZ; DBS-S were crossed with males FRT40A to obtain +/+ genetic mosaics in a M/+ genetic background, thus triggering cell competition. Progeny of this cross were heat-shocked at 48-72h after egg laying (AEL) 37°C for 1 h. The larvae were dissected before puparation to perform the immunostainings in Figure 1E.

### Temperature shift experiments

After 12h of egg laying at 25°C, crossed flies were transferred into new tubes. Eggs laid were kept at 22°C for three days in order to prevent lethality at early developmental stages. Hatched larvae were then transferred to 25°C until dissection before puparium formation. The genotypes of the flies used in these experiments were:

w; *UAS-Hep.Act, Tubulin-DBS-S /TubG80*^*ts*^; *Hedgehog-Gal4/UAS-P35 (Figure 3C)*

w; *UAS-Hid, Tubulin-DBS-S /TubG80*^*ts*^; *Hedgehog-Gal4/UAS-P35 (Figure 3D)*

### Immunohistochemistry

Immunostainings and washes of imaginal discs and S2 cells were performed according to standard protocols (fixing in PBS 4% paraformaldehyde, washing in PBT 0.3% (0.3% Triton X-100 in PBS)). Adult *Drosophila* intestines were dissected in ice-cold PBS. To fix, wash solution (0.7% NaCl, 0.05% Triton X-100) was heated to approximately 90°C and the intestines submerged into this solution for 6 seconds. Following this the intestines were rapidly cooled through submersion in ice cold wash solution. Three rapid washes in PBT (0.3%) preceded blocking in 1% BSA PBT (0.3%) for at least one hour. For imaginal disc immunostaining, primary and secondary antibody incubations were made at room temperature for three hours. For gut immunostaining, overnight incubation at 4°C with the primary antibodies and two hours at room temperature with the secondary antibodies was necessary. Primary antibodies used in our experiments were: anti-cleaved Caspase3 (1:100; Cell Signalling 9661); anti-P35 (1:500; Imgenex, IMG5740); rabbit anti-HA (1:1000; Cell signaling C29F4); mouse anti-betaGal (1:500; Promega Z378B), chicken Anti-betaGal (1:200, Abcam AB9361). Anti-Armadillo (1:50, Hybridoma Bank N2 7A1 armadillo-c); Anti-Prospero (1:20, Hybridoma Bank MR1A). Conjugated secondary antibodies (Molecular Probes) were diluted in 0.3% PBT and used in a final concentration (1:200): anti-mouse Alexa 555, anti-mouse Alexa 488, anti-mouse Alexa 647, anti-rabbit Alexa 555, anti-rabbit Alexa 633. Following incubation in secondary antibodies, samples were washed in PBT several times during 60 minutes and mounted in Vectashield with DAPI (1:1000; Thermo Scientific 62248).

### Irradiation protocol

Larvae were exposed to a source of γ-ray during the irradiation experiments. Time exposure was adjusted for administrating a dose of 1500 rads to the samples.

### Western blot

Protein extracts were obtained from either non-irradiated or irradiated larvae (4h post-treatment). We then followed standard protocols for processing samples and immunoblotting the membranes. We used anti-GFP (SIGMA-11814460001; 1:3000) as primary antibody in these experiments.

### Cell culture and transfection of S2 and S2-R+ cells

S2 and S2-R+ cells were grown in Gibco Schneider’s medium supplemented with 1% L-glutamine, 10% fetal calf serum and antibiotics (1% penicillin/streptavidin). In all of the experiments, 1 microgram of plasmid was transfected using Effectene (Qiagen) reagent, following manufacturer’s instructions. Induced apoptosis was achieved after exposing the cells for 10 minutes to UV light. Upon UV light treatment, cells were then fixed and immunolabelled at different time points (1h:30’ and 4h). In most experiments, cells were grown on normal coverslips placed in 6-well plates; however, coverslips coated with poly-lysine (SIGMA P0425-72EA) were used to facilitate the visualization of cell projections and active actin in Figure S1E.

### Imaging of fixed and live samples

Fluorescent imaging of fixed and live imaginal discs was performed with a Leica SP8 laser-scanning confocal microscope using LAS AF software. *Drosophila* intestinal fixed samples were imaged using the Olympus Fluoview FV1200 and associated software. Z-stacks were taken with a 40X objective at intervals along the apical-basal axis that ensured adequate resolution in Z (step size 0.5-1.5-µm). Acquired images were then processed using ImageJ and Adobe Photoshop CS6. Culture conditions and live recording of imaginal discs was performed as described by Mao and collaborators (Mao et al., 2011). 90 minutes after irradiation, wing imaginal discs were filmed for 40-60 minutes. 4-6 focal planes covering the subapical region of the wing disc epithelia were projected using ImageJ in order to obtain full representation of the nucleus and actin projections from single cells (Movie 1). The reslice function of Image J. was used to generate cross-sectional movies (Movies 2-4); again 4-6 focal planes were merged in order to obtain full representation of the nucleus and actin projections from single cells.

Heatmaps in Figure 5 were produced using Fiji (ImageJ) software. Original images were acquired with the same orientation and magnification. The enriched area in stem cells of the R5 segment was used as reference to align the guts. On average, 20 focal planes per gut, containing all the information from the apical to the middle sections of a Z-stack, were projected using the Average intensity function. Subsequently, all the channels not containing information related with reporter’s activation were discarded. The projected reporter channel images were combined into a new single Z-stack of projections, before applying an average intensity projection again. Following this we applied the 16 colour LUT.

Dot Plots in Supplementary Figure 4 were produced using Fiji (ImageJ) software. Dot plots were created by combining all of the channels (RGB) into one channel. Following this, a new black background channel was inserted. A composite image was selected and the pencil tool used to mark (white) the location of presumptive progenitors cells onto the black channel. The nuclear size and armadillo/prospero immunostaining was used to assign the cell fate to different intestinal cells. A projection of this channel was created and for quantification purposes the dots separated on the projected image. Three equal sections of the resultant picture were counted sequentially using the Analyse particle function and the percentage of enteroblasts in each location calculated.

4D microscopy of the abdominal epidermis was performed with a Leica SP8 confocal microscope at 25±1°C. Pupae were staged according to (Bainbridge and Bownes, 1981) and then were dissected and filmed as described in (Seijo-Barandiaran et al., 2015). A step size of 2.5 µm in Z was used during the filming to image the samples. Samples were imaged using a time interval range of 120s to 180s, depending on the experiment.

## ACKNOWLEDGEMENTS

Thanks for providing flies and reagents to; Paul D. Friesen (University of Wisconsin-Madison; cDNA of Drice containing the mutation C 211> A); Elke Ober (cDNA of lifeAct-Ruby); Jose Felix de Celis (*spalt*-Gal4; Centro de Biología Molecular); Andreas Bergmann (GUS-dp53, GMR.P53 lines), Antonio Baonza (*hs-flipase*; *FRT40 Minute arm-LacZ)*. Thanks also to Jordan Raff, Antonio Baonza and caspaselab members (https://www.caspaselab.com) for the critical reading of the manuscript and valuable suggestions. Bestgene and Genewiz for generating transgenic flies and synthetic DNA constructs, respectively. This work has been supported by Cancer Research UK C49979/A17516 and the John Fell Fund from the University of Oxford 162/001. L.A.Baena-Lopez is a CRUK Career Development Fellow (C49979/A17516) and an Oriel College Hayward Fellow. L. Arthurton is a PhD student supported by the Edward Penley Abraham Research Fund. M. Bischoff is supported by BBSRC (BB/M021084/1). J-P.V. and C.A are supported by an ERC grant WNTEXPORT; 294523 and the MRC MRC U117584268.

## CONTRIBUTIONS

L.A.B.-L. was responsible for the conception of the work, text of the manuscript, figure preparation and most of the experimental work, including the molecular cloning. During the initial stages of the project R. McGregor as undergraduate rotational student assisted L.A.B-L with the experimental work (Figures 1, 3 and S1). L.A. has performed the experiments and data analysis related to the intestinal system (Figures 5 and S4). M. B completed the live imaging in pupal *Drosophila* abdomen (Figure S2, Movie 6 and Movie 7). C.A. generated the starting plasmid to build the 2A constructs; he has also provided critical molecular cloning advice. J.P.V. provided critical guidance at the initial stages of the project; part of the work was also performed during the stay of L.A. B-L in his laboratory. All co-authors have provided useful criticisms and commented on the manuscript before submission.

## REFERENCES

Abrams, J.M., White, M.A., 2004. Coordination of cell death and the cell cycle: linking proliferation to death through private and communal couplers. Curr Opin Cell Biol 16, 634–638.

Aram, L., Yacobi-Sharon, K., Arama, E., 2017. CDPs: caspase-dependent non-lethal cellular processes. Cell Death Differ 24, 1307–1310.

Baena-Lopez, L.A., Arthurton, L., Xu, D.C., Galasso, A., 2017. Non-apoptotic Caspase regulation of stem cell properties. Semin Cell Dev Biol.

Bainbridge, S.P., Bownes, M., 1981. Staging the metamorphosis of Drosophila melanogaster. J Embryol Exp Morphol 66, 57–80.

Bardet, P.L., Kolahgar, G., Mynett, A., Miguel-Aliaga, I., Briscoe, J., Meier, P., Vincent, J.P., 2008. A fluorescent reporter of caspase activity for live imaging. Proc Natl Acad Sci U S A 105, 13901–13905.

Bischoff, M., Cseresnyes, Z., 2009. Cell rearrangements, cell divisions and cell death in a migrating epithelial sheet in the abdomen of Drosophila. Development 136, 2403–2411.

Buchon, N., Osman, D., David, F.P., Fang, H.Y., Boquete, J.P., Deplancke, B., Lemaitre, B., 2013. Morphological and molecular characterization of adult midgut compartmentalization in Drosophila. Cell Rep 3, 1725–1738.

Burgon, P.G., Megeney, L.A., 2017. Caspase signaling, a conserved inductive cue for metazoan cell differentiation. Semin Cell Dev Biol.

Ding, A.X., Sun, G., Argaw, Y.G., Wong, J.O., Easwaran, S., Montell, D.J., 2016. CasExpress reveals widespread and diverse patterns of cell survival of caspase-3 activation during development in vivo. Elife 5.

Ellis, H.M., Horvitz, H.R., 1986. Genetic control of programmed cell death in the nematode C. elegans. Cell 44, 817–829.

Florentin, A., Arama, E., 2012. Caspase levels and execution efficiencies determine the apoptotic potential of the cell. J Cell Biol 196, 513–527.

Fogarty, C.E., Bergmann, A., 2017. Killers creating new life: caspases drive apoptosis-induced proliferation in tissue repair and disease. Cell Death Differ 24, 1390–1400.

Gudipaty, S.A., Rosenblatt, J., 2017. Epithelial cell extrusion: Pathways and pathologies. Semin Cell Dev Biol 67, 132–140.

Hay, B.A., Wolff, T., Rubin, G.M., 1994. Expression of baculovirus P35 prevents cell death in Drosophila. Development 120, 2121–2129.

Hollville, E., Deshmukh, M., 2017. Physiological functions of non-apoptotic caspase activity in the nervous system. Semin Cell Dev Biol.

Kanuka, H., Kuranaga, E., Takemoto, K., Hiratou, T., Okano, H., Miura, M., 2005. Drosophila caspase transduces Shaggy/GSK-3beta kinase activity in neural precursor development. EMBO J 24, 3793–3806.

Kawamoto, Y., Nakajima, Y.I., Kuranaga, E., 2016. Apoptosis in Cellular Society: Communication between Apoptotic Cells and Their Neighbors. Int J Mol Sci 17.

Kerr, J.F., Wyllie, A.H., Currie, A.R., 1972. Apoptosis: a basic biological phenomenon with wide-ranging implications in tissue kinetics. Br J Cancer 26, 239–257.

Kondo, S., Senoo-Matsuda, N., Hiromi, Y., Miura, M., 2006. DRONC coordinates cell death and compensatory proliferation. Mol Cell Biol 26, 7258–7268.

LaCount, D.J., Hanson, S.F., Schneider, C.L., Friesen, P.D., 2000. Caspase inhibitor P35 and inhibitor of apoptosis Op-IAP block in vivo proteolytic activation of an effector caspase at different steps. J Biol Chem 275, 15657–15664.

Lannan, E., Vandergaast, R., Friesen, P.D., 2007. Baculovirus caspase inhibitors P49 and P35 block virus-induced apoptosis downstream of effector caspase DrICE activation in Drosophila melanogaster cells. J Virol 81, 9319–9330.

Levayer, R., Dupont, C., Moreno, E., 2016. Tissue Crowding Induces Caspase-Dependent Competition for Space. Curr Biol 26, 670–677.

Mao, Y., Tournier, A.L., Bates, P.A., Gale, J.E., Tapon, N., Thompson, B.J., 2011. Planar polarization of the atypical myosin Dachs orients cell divisions in Drosophila. Genes Dev 25, 131–136.

Marinari, E., Mehonic, A., Curran, S., Gale, J., Duke, T., Baum, B., 2012. Live-cell delamination counterbalances epithelial growth to limit tissue overcrowding. Nature 484, 542–545.

McIlwain, D.R., Berger, T., Mak, T.W., 2013. Caspase functions in cell death and disease. Cold Spring Harbor perspectives in biology 5, a008656.

Meyer, E.J., Ikmi, A., Gibson, M.C., 2011. Interkinetic nuclear migration is a broadly conserved feature of cell division in pseudostratified epithelia. Curr Biol 21, 485–491.

Miura, M., 2012. Apoptotic and nonapoptotic caspase functions in animal development. Cold Spring Harb Perspect Biol 4.

Moreno, E., Basler, K., Morata, G., 2002. Cells compete for decapentaplegic survival factor to prevent apoptosis in Drosophila wing development. Nature 416, 755–759.

Mukherjee, A., Williams, D.W., 2017. More alive than dead: non-apoptotic roles for caspases in neuronal development, plasticity and disease. Cell Death Differ 24, 1411– 1421.

Napoletano, F., Gibert, B., Yacobi-Sharon, K., Vincent, S., Favrot, C., Mehlen, P., Girard, V., Teil, M., Chatelain, G., Walter, L., Arama, E., Mollereau, B., 2017. p53-dependent programmed necrosis controls germ cell homeostasis during spermatogenesis. PLoS Genet 13, e1007024.

Ninov, N., Chiarelli, D.A., Martin-Blanco, E., 2007. Extrinsic and intrinsic mechanisms directing epithelial cell sheet replacement during Drosophila metamorphosis. Development 134, 367–379.

Ninov, N., Martin-Blanco, E., 2007. Live imaging of epidermal morphogenesis during the development of the adult abdominal epidermis of Drosophila. Nat Protoc 2, 3074–3080.

Ohsawa, S., Vaughen, J., Igaki, T., 2018. Cell Extrusion: A Stress-Responsive Force for Good or Evil in Epithelial Homeostasis. Dev Cell 44, 284–296.

Ouyang, Y., Petritsch, C., Wen, H., Jan, L., Jan, Y.N., Lu, B., 2011. Dronc caspase exerts a non-apoptotic function to restrain phospho-Numb-induced ectopic neuroblast formation in Drosophila. Development 138, 2185–2196.

Perez-Garijo, A., 2017. When dying is not the end: Apoptotic caspases as drivers of proliferation. Semin Cell Dev Biol.

Perez-Garijo, A., Martin, F.A., Struhl, G., Morata, G., 2005. Dpp signaling and the induction of neoplastic tumors by caspase-inhibited apoptotic cells in Drosophila. Proc Natl Acad Sci U S A 102, 17664–17669.

Perez-Garijo, A., Shlevkov, E., Morata, G., 2009. The role of Dpp and Wg in compensatory proliferation and in the formation of hyperplastic overgrowths caused by apoptotic cells in the Drosophila wing disc. Development.

Potter, C.J., Tasic, B., Russler, E.V., Liang, L., Luo, L., 2010. The Q system: a repressible binary system for transgene expression, lineage tracing, and mosaic analysis. Cell 141, 536–548.

Prokhorova, E.A., Zamaraev, A.V., Kopeina, G.S., Zhivotovsky, B., Lavrik, I.N., 2015. Role of the nucleus in apoptosis: signaling and execution. Cell Mol Life Sci 72, 4593–4612.

Ramirez, M.L.G., Salvesen, G.S., 2018. A primer on caspase mechanisms. Semin Cell Dev Biol.

Schott, S., Ambrosini, A., Barbaste, A., Benassayag, C., Gracia, M., Proag, A., Rayer, M., Monier, B., Suzanne, M., 2017. A fluorescent toolkit for spatiotemporal tracking of apoptotic cells in living Drosophila tissues. Development 144, 3840–3846.

Seijo-Barandiaran, I., Guerrero, I., Bischoff, M., 2015. In Vivo Imaging of Hedgehog Transport in Drosophila Epithelia. Methods Mol Biol 1322, 9–18.

Songane, M., Khair, M., Saleh, M., 2018. An updated view on the functions of caspases in inflammation and immunity. Semin Cell Dev Biol.

Takemoto, K., Nagai, T., Miyawaki, A., Miura, M., 2003. Spatio-temporal activation of caspase revealed by indicator that is insensitive to environmental effects. J Cell Biol 160, 235–243.

Tang, H.L., Tang, H.M., Fung, M.C., Hardwick, J.M., 2015. In vivo CaspaseTracker biosensor system for detecting anastasis and non-apoptotic caspase activity. Sci Rep 5, 9015.

To, T.L., Piggott, B.J., Makhijani, K., Yu, D., Jan, Y.N., Shu, X., 2015. Rationally designed fluorogenic protease reporter visualizes spatiotemporal dynamics of apoptosis in vivo. Proc Natl Acad Sci U S A 112, 3338–3343.

Wells, B.S., Yoshida, E., Johnston, L.A., 2006. Compensatory proliferation in Drosophila imaginal discs requires Dronc-dependent p53 activity. Curr Biol 16, 1606–1615.

Zaret, K.S., 2014. Genome reactivation after the silence in mitosis: recapitulating mechanisms of development? Dev Cell 29, 132–134.

